# EMPAC: A Multimodal Dataset for Bridging Affective and Cognitive Empathy

**DOI:** 10.64898/2026.05.11.724205

**Authors:** Airi Ota, Shiro Kumano, Aiko Murata, Ai Nakane, Shinya Shimizu

## Abstract

Empathy, a key element of social interaction, involves both cognitive and affective processes and is commonly investigated through measures such as empathic accuracy and affective physiological synchrony. While physiological synchrony offers a continuous measure of affective processes, empathic accuracy typically relies on discrete self-reports, leaving their temporal relationship largely unexplored. Advancing this line of research requires datasets that integrate time-continuous self-reports with physiological signals, yet such datasets—particularly those focusing on the empathizee—remain limited. To fill this gap, we present EMPAC (Empathy Measurement: Physiological, Affective, and Cognitive), a multimodal dataset constructed. To create empathy-eliciting stimuli, professional actors performed emotionally intense, pseudo-autobiographical narratives while their physiological signals (e.g., ECG, EDA) and continuous self-reported emotional states were recorded. We then conducted two observer experiments using these video recordings. In Experiment 1, to validate the stimuli as empathy-eliciting materials, observers continuously rated emotional intensity without being informed of the specific emotion portrayed, following the protocol of previous studies on time-series empathic accuracy. Yet this approach sometimes revealed a gap between the emotion category portrayed by the target and that perceived by the observers. In Experiment 2, we introduced a revised procedure in which the target emotion category was disclosed prior to viewing, revealing that specifying the target emotion led to a different relationship between individual empathy traits and empathic accuracy than observed in Experiment 1. EMPAC thus provides a rich, temporally aligned resource for investigating empathy dynamics in naturalistic settings and for evaluating methodological variations in empathic accuracy paradigms.

## 1 INTRODUCTION

Empathy, commonly conceptualized as the interplay of cognitive and affective responses arising from observing another individual’s emotional experiences, is crucial for social connection and effective communication. These cognitive and affective empathic responses vary widely among individuals, influenced by factors such as neurodevelopmental conditions and personality traits (Decety and Jackson, 2004). For people who struggle with empathy, the development of accurate empathy recognition technologies is highly desirable. However, progress in this area is hindered by the limited availability of comprehensive datasets that capture empathy’s multifaceted nature, spanning both its cognitive and affective dimensions.

Cognitive empathy refers to an individual’s capacity to understand and accurately identify the feelings and perspectives of others. It involves recognizing and interpreting emotional expressions, thoughts, and intentions (Ickes, 1993). A key metric in this domain is empathic accuracy (EA), which quantifies how closely an observer’s (the empathizer’s) judgments of another person’s (the empathizee’s) emotional state align with that person’s self-reported experiences during an interaction. Two types of paradigms have been developed for assessing EA. The *Ickes paradigm*, the most widely used approach, involves dyadic interaction or standard stimulus approaches, in which perceivers infer the target’s thoughts and feelings at specific moments and accuracy is scored by comparing these inferences with the target’s own reports (Ickes, 1993). In contrast, the *Zaki paradigm*, a newer method increasingly adopted in social neuroscience research, employs continuous affect rating procedures, where both targets and perceivers provide dynamic reports of affective intensity while viewing the same video, and empathic accuracy is quantified by correlating their time series (Zaki et al., 2008). Recent studies underscore the value of such continuous measures for accurately reflecting the fluid, moment-to-moment nature of cognitive empathy (Mackes et al., 2018; McKenzie et al., 2022).

Affective empathy, by contrast, involves sharing or mirroring the emotional experiences of others. It centers on the empathizer’s affective responses, often underpinned by mirror systems, which can lead to emotional synchronization with the empathizee (Gallese, 2001). Studies have documented synchronized neural and physiological patterns—such as electroencephalograms (EEG), Electrocardiograms (ECG), electrodermal activity (EDA), and blood volume pulses (BVP)—during expressive emotional interactions (Anders et al., 2011). Such physiological synchronization has been linked to trait sociality, suggesting that affective empathic states may be inferred through these shared physiological signals (Murata et al., 2020). Consequently, measuring affective empathy frequently entails analyzing physiological data to detect parallel emotional responses across interacting individuals.

Despite the importance of examining cognitive and affective empathy in tandem, few publicly available datasets capture both aspects simultaneously. This shortfall partly reflects the absence of established experimental paradigms capable of collecting key data from both empathizees and empathizers. One challenge lies in obtaining empathizees’ physiological responses that reliably reflect target affective states (e.g., anger or surprise) while also featuring pronounced affective expressions—such as distinct facial and vocal cues—needed by empathizers for accurate cognitive empathy assessment. At the same time, recording robust physiological signals from empathizers is essential for evaluating affective empathy, but these data can be noisy, complicating the measurement of interpersonal synchrony. This is because empathizers’ autonomic responses are elicited only indirectly—via perception and inference from the empathizee’s expressed behavior—whereas empathizees’ physiology is more proximally coupled to the affective state they enact (e.g., through recall or simulation); consequently, unless the empathizee’s affect is clearly and normatively expressed (as is more likely with professionally trained actors), observers’ psychophysiological responses tend to be attenuated and variable, reducing apparent synchrony (Palumbo et al., 2017; Zaki et al., 2008). Additionally, it is ethically and logistically difficult to elicit strong emotions in experimentally naive participants (acting as empathizees) and ensure they communicate these states effectively to others. As a result, existing datasets seldom satisfy all these requirements, as summarized in Table 1, and thus fail to offer the detailed insights required for in-depth empathy research.

**Table 1.**
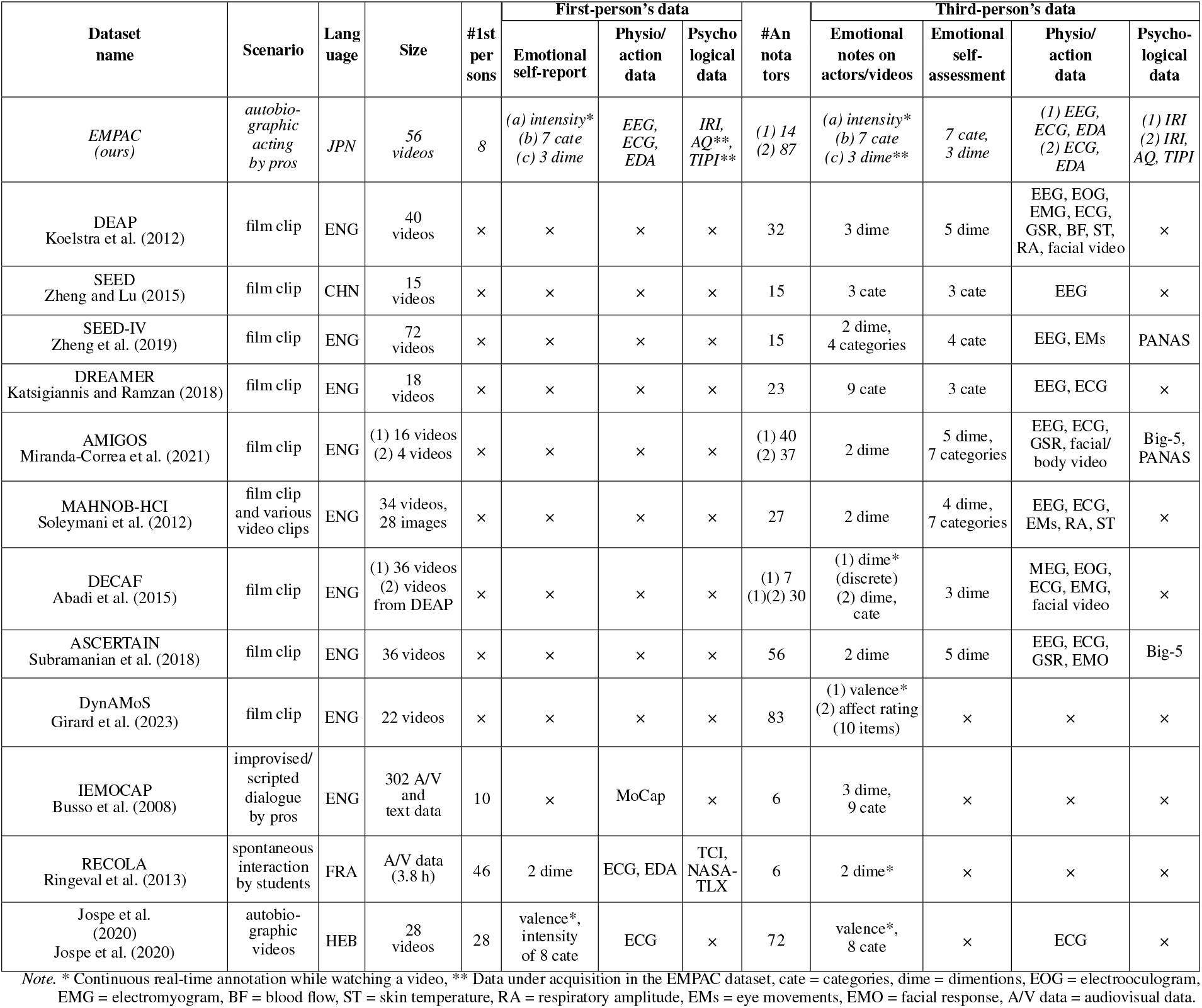
EXISTING EMOTIONAL STIMULUS DATASET AND OUR DATASET.

To help overcome these challenges and enable the joint examination of cognitive and affective empathy, we introduce a new dataset called EMPAC (Empathy Measurement: Physiological, Affective, and Cognitive) dataset. This resource is designed to support comprehensive empathy research by capturing both high-quality physiological data and time-continuous emotional annotations from empathizees and empathizers. Specifically, it comprises 24 pseudo-autobiographical videos featuring eight professional actors (as empathizees), plus corresponding data from 40 observers (as empathizers). To facilitate in-depth investigations, the dataset includes:

- Physiological Responses: ECG and EDA from both empathizees and empathizers, enabling the study of physiological synchronization.
- Time-Continuous Emotion Intensity Annotations: Both actors’ self-reports and observers’ intensity ratings, supporting EA analysis over time.
- Controlled Emotional Stimuli: Professionally acted emotional expressions employing ethically sound elicitation methods, ensuring pronounced affective cues for reliable observation.

We combined autobiographical recall strategies with the acting methods outlined by Hodge (Hodge, 2000), guiding professional actors to draw on personal memories but present them in pseudo-biographical form (Siedlecka and Denson, 2019). Because these performances do not recreate the actors’ actual life events, we alleviate concerns about disclosing sensitive personal experiences of the empathizees, yet still obtain high-intensity expressions. Such vivid portrayals, in turn, promote strong emotional engagement from naive empathizers, enabling accurate perception of the displayed emotions (Zaki et al., 2009). Drawing on the century-long evolution of actor training described by Hodge (Hodge, 2000), professional performers are particularly adept at harnessing techniques from Stanislavski-based systems—such as structured imagination, emotional memory, and disciplined physicality—to evoke intense emotional states in a controlled manner. This allows us to bypass the ethical and practical difficulties of eliciting comparable reactions from naive participants, while ensuring the authenticity of the emotional stimuli. Although few studies have collected physiological data from professional actors (Malcolm, 2012), prior findings indicate that subjective arousal closely aligns with physiological activation (Kreibig, 2010a). In our dataset, we likewise observed robust physiological changes among the actors, underscoring the validity of this approach to capturing both affective intensity and consistency in emotional expression.

Our dataset fills a critical gap in empathy research by capturing both the cognitive dimension—through empathizers’ accurate interpretation of empathizees’ emotions—and the affective dimension—through physiological synchronization between empathizees and empathizers. This integrated perspective enables a more nuanced investigation of empathy, facilitating the study of both components in tandem. By offering a rich dataset of synchronized physiological signals and emotional annotations, we aim to drive advances in empathy recognition technologies and foster a deeper understanding of empathic processes.

Although our dataset includes both subjective annotations and physiological data, the present study primarily validates the video stimuli using actors’ self-reports, their physiological responses, and external observers’ annotations. In this validation, we conducted two experiments in which external observers watched the videos and continuously rated the actors’ emotional intensity. We then computed EA scores by correlating the actors’ time-continuous self-reports with the observers’ ratings, consistent with established methods (McKenzie et al., 2022). For physiological measurements, we employed heart rate (HR) variability analysis, ensuring robust and reliable data collection.

In our initial experiment, aligned with prior studies (Mackes et al., 2018) that did not define specific target emotions for time-continuous intensity ratings, we found a positive correlation between EA scores and the perspective-taking trait of the observers. Notably, we obtained similar results even though the original study was conducted in French whereas our experiment was conducted in Japanese, illustrating the robustness of the EA paradigm across different languages and cultures. However, as acknowledged in that earlier work, emotional intensity judgment relies on brain regions distinct from those involved in categorical or dimensional judgments: the amygdala processes intensity, while the orbitofrontal cortex is associated with valence (Lewis et al., 2007). When we subsequently specified the target emotion during continuous ratings, the correlation with perspective-taking disappeared, underscoring the importance of clarity in empathy research and suggesting a need to explore multiple approaches to understanding empathic processes.

To facilitate further research in this domain, we provide access to the EMPAC dataset—comprising time-continuous emotion ratings, ECG and EDA recordings from both professional actors and observers—for academic use upon reasonable request and subject to institutional review. In compliance with local ethical and legal requirements, data access is currently limited to researchers affiliated with institutions in Japan. This restriction reflects regulatory obligations rather than technical limitations, and ensures that the dataset is shared in a manner consistent with the highest ethical standards. Access is managed through a dedicated contact point and requires agreement with our institution’s data use policy. At present, availability is focused on the video stimuli and actor-related data described in this study, with plans to expand access as additional components of the dataset are curated. While current access is restricted to Japanese institutions, we are actively exploring pathways for broader, international availability in accordance with evolving ethical and legal frameworks. EMPAC is intended to serve as a long-term research resource, supporting advancements in empathy modeling, affective computing, and mental health applications.

## 2 EXISTING EMPATHIC VIDEO STIMULI

Affective empathy involves generating emotional responses similar to the target’s, measured by neural or autonomic responses like heart rate and skin conductance (Prochazkova and Kret, 2017). Cognitive empathy entails understanding the target’s emotions, measured by how accurately participants appraise others’ emotions using subjective reports (Zaki and Ochsner, 2011). Given this and the validity of autobiographical recall methods, we consider our method sound. We showcase in Table 1 that our dataset uniquely measures both types of empathy.

The exploration of human emotions through visual stimuli has significantly advanced with the aid of various emotion-focused datasets such as DEAP (Koelstra et al., 2012), SEED (Zheng and Lu, 2015), SEED-IV (Zheng et al., 2019), DREAMER (Katsigiannis and Ramzan, 2018), AMIGOS (Miranda-Correa et al., 2021), MAHNOB-HCI (Soleymani et al., 2012), DECAF (Abadi et al., 2015), ASCERTAIN (Subramanian et al., 2018), DynAMoS (Girard et al., 2023), IEMOCAP (Busso et al., 2008), and RECOLA (Ringeval et al., 2013). As summarized in Table 1, These datasets, spanning a range of scenarios from commercial music videos to spontaneous interactions, delve deeply into the complexities of emotional responses, offering invaluable insights for the broader field of emotion research. The diversity of these datasets in structure and data types lays the groundwork for understanding the nuances of emotional experiences. For example, while DEAP and DREAMER focus on film clips to examine emotional responses, IEMOCAP and RECOLA provide insights into more spontaneous and nuanced emotional exchanges. This variance in data collection and analysis methods forms a crucial bridge to the exploration of empathetic video stimuli, allowing a comprehensive examination of emotional expressions and interactions.

When examining empathic accuracy literature within psychology and neuroscience, where datasets are rarely made publicly available, research is often categorized into film clips, autobiographical, and dialogical formats, paralleling the primary datasets. Film clips, akin to those used in DEAP and DREAMER, provide rich emotional expressions but can pose challenges in clearly identifying empathy targets. Autobiographical formats, reflecting the depth of self-reported experiences as seen in datasets like AMIGOS, offer detailed insights into personal emotional journeys, though they may lack comprehensive physiological data integration. Similarly, autobiographical format known as empathic accuracy tasks (Mackes et al., 2018; McKenzie et al., 2022) and dialogical formats as demonstrated by the annotations in IEMOCAP, emphasize the importance of time-continuous annotations in capturing the subtleties of emotional dynamics. This interplay between primary emotion-focused datasets and empathic video stimuli highlight the ongoing challenges and opportunities in the field, emphasizing the need for a holistic approach to deepen our understanding of empathy and emotional responses.

In this context, our dataset, EMPAC, offers a distinctive contribution. Focusing on autobiographic acting by professional actors, EMPAC provides a unique blend of first-person and third-person emotional data. This dataset not only captures the actors’ self-reported emotional experiences but also includes physiological and psychological measurements, enriching the understanding of emotional expression and perception. By offering a detailed account of both the actors’ and observers’ perspectives on a range of emotions, EMPAC stands out as a valuable resource for exploring the intricacies of emotional experiences, complementing the broader array of datasets in the field and advancing our comprehension of empathy in a cross-cultural context.

## 3 OUR VIDEO STIMULI (ACTORS’ DATA)

In this study, in response to the challenges with existing empathetic video stimuli, we constructed a new set of pseudo-autobiographical video stimuli featuring characteristics outlined in Table 1. The procedure fundamentally adhered to the existing EA protocol (McKenzie et al., 2022), commonly used in various fMRI studies to assess empathic accuracy and capture the neurological basis of empathy (Mackes et al., 2018; McKenzie et al., 2022). Emotional labels were set based on existing stimuli, including both emotion category labels and continuous emotion intensity labels. Based on Ekman and Friesen’s six basic emotions (Ekman and Friesen, 1976), we predefined seven emotions per video—anger, disgust, fear, joy, sadness, surprise, plus neutral. Post-filming, the actors themselves time-continuously assigned emotion intensity on a nine-point scale. Note that for the actors, the target emotion they rated was obvious, as each actor was instructed to express only the predetermined emotion category.

### 3.1 Actors

To create the video stimuli, we engaged eight professional Japanese actors, ranging in age from their 20s to 60s, all skilled in eliciting and expressing emotions. Recruitment was conducted from March 4 to March 8, 2022, in compliance with ethical requirements. On the day of data collection, the experimenter explained the study orally using a written information sheet and consent form, read the contents aloud, and obtained written informed consent from all participants prior to the start of the experimental procedure.

### 3.2 Procedure

#### 3.2.1 Filming

Each actor narrated pseudo-autobiographical emotional events in videos lasting 1–3 minutes. These stories were adapted from real-life experiences but modified to enhance emotional intensity while protecting privacy. To limit workload, each actor performed three distinct episodes selected from the six non-neutral emotion categories (joy, sadness, anger, fear, disgust, surprise), resulting in four actors assigned to each category. In addition, all eight actors recorded one neutral video in which they described the layout of their home without intentionally evoking any emotion.

Each emotional episode was filmed under two conditions: (1) with-expression condition, in which actors expressed emotions *overtly* through facial expressions and vocal tone; and (2) without-expression condition, in which actors maintained a neutral tone and facial expression, *suppressing* emotional expression during speech.

The without-expression condition was designed to specifically target emotion regulation—particularly emotion suppression, an area underexplored in affective computing—and thus distinguishes our dataset from previous ones. In both conditions, prior to filming each emotional episode, actors were instructed to *internally* evoke the target emotion by recalling the original experience using affective memory techniques inspired by Strasberg and Stanislavski. To verify emotional arousal, filming began only when the actor’s heart rate, relative to the baseline established during neutral recording, had increased for at least 5 seconds or remained elevated above baseline for more than 5 seconds. This procedure ensured that each actor’s emotional response was genuinely elicited.

In total, this procedure yielded 56 videos (4 *×* 2 *×* 6 + 8): four actors *×* two expression conditions (with/without expression) *×* six emotions, plus one neutral video per actor. Although the neutral and without-expression videos are not the primary focus of this study, all conditions will be made publicly available.

#### 3.2.2 Emotional intensity annotation by actors

After filming, the actors watched the videos in which they performed and time-continuously rated their emotional intensity on the same day using a 9-point scale (from 1, ‘no emotion’, to 9, ‘very strong emotion’). We strictly adhered to the procedure outlined in the referenced literature (Mackes et al., 2018), although dimension-continuous annotation, such as the visual analogue scale, is also used for time-continuous annotation.

#### 3.2.3 Physiological measurement of actors

Physiological signals were recorded from the actors during both filming and subsequent annotation. ECG was employed to capture heart rate as an index of autonomic nervous system activity. EDA and EEG were also collected, although they were not analyzed in the present study. ECG and EDA were measured using the Intercross-415 (Intercross Corp.), and EEG was recorded with the VIE ZONE (Vie Style Inc.) earphone-type brainwave monitor, selected for its ease of use.

#### 3.2.4 Personality trait measurement of actors

Finally, to assess actors’ dispositional empathy, they completed the Japanese version of the Interpersonal Reactivity Index (Davis, 1980); IRI-J (Himichi et al., 2017). This instrument comprises four subscales: Fantasy (FS), reflecting affective empathy through engagement with fictional characters; Empathic Concern (EC), indicating feelings of warmth and concern for others; Perspective Taking (PT), representing the cognitive aspect of empathy by adopting others’ viewpoints; and Personal Distress (PD), measuring self-oriented feelings of anxiety and discomfort in tense interpersonal situations.

### 3.3 Analysis for validating actors’ affective evocation on heart rate

We examined whether the actors elicited the intended target emotional responses during both acting and self-annotation across all 56 videos. R-R intervals (RRI) and heart rate (HR) were first calculated from the ECG data to obtain mean HR for each condition, using MATLAB 9.14.0 and the Signal Processing Toolbox (MathWorks, Inc.). We then conducted a linear mixed model (LMM) analysis of mean HR with the lmer function in the lmerTest package in R 4.4.0 (R Core Team, 2024), in order to assess changes in emotional responses at the physiological level. In the LMM, condition (neutral videos, videos with emotional expression, and videos without emotional expression) and task (acting vs. annotation) were specified as fixed effects, and actor ID was specified as a random effect to account for individual differences in heart rate responses.

To further investigate temporal changes in emotional arousal within each video, average HRs were calculated separately for the first and second halves of each video and compared across the 24 videos with emotional expressions, during both the filming and annotation phases. Pair plots and statistical analyses were conducted in R 4.4.0. Shapiro–Wilk normality tests indicated that the data did not follow a normal distribution; therefore, Wilcoxon signed-rank tests were applied for subsequent comparisons.

### 3.4 Results

#### 3.4.1 Qualitative evaluation of actors’ annotations and heart rate

Figure 2 illustrates examples in which actors’ HR during emotional acting closely corresponded to their own continuous intensity annotations.

**Figure 1.**
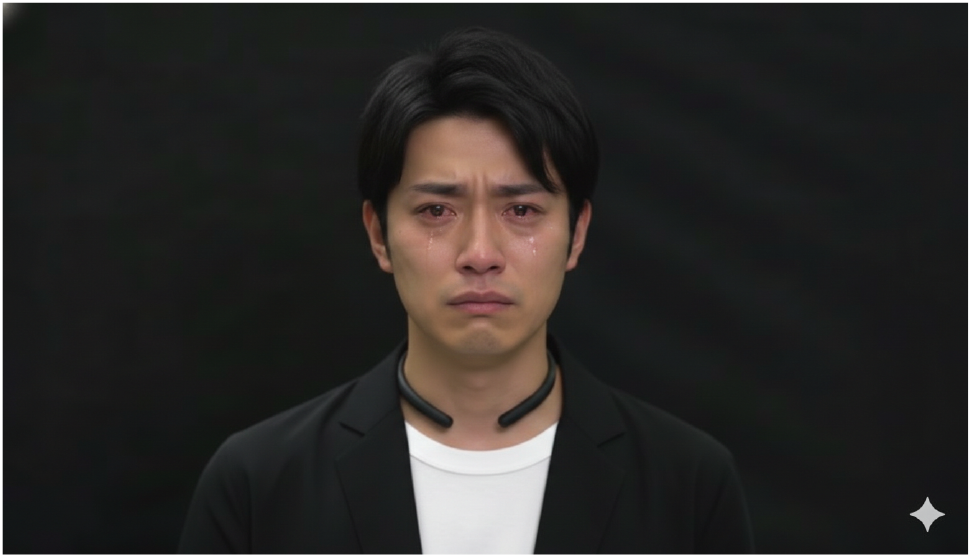
Example stimulus video of “sadness”. The image shown here was generated using generative AI for illustrative purposes.

**Figure 2.**
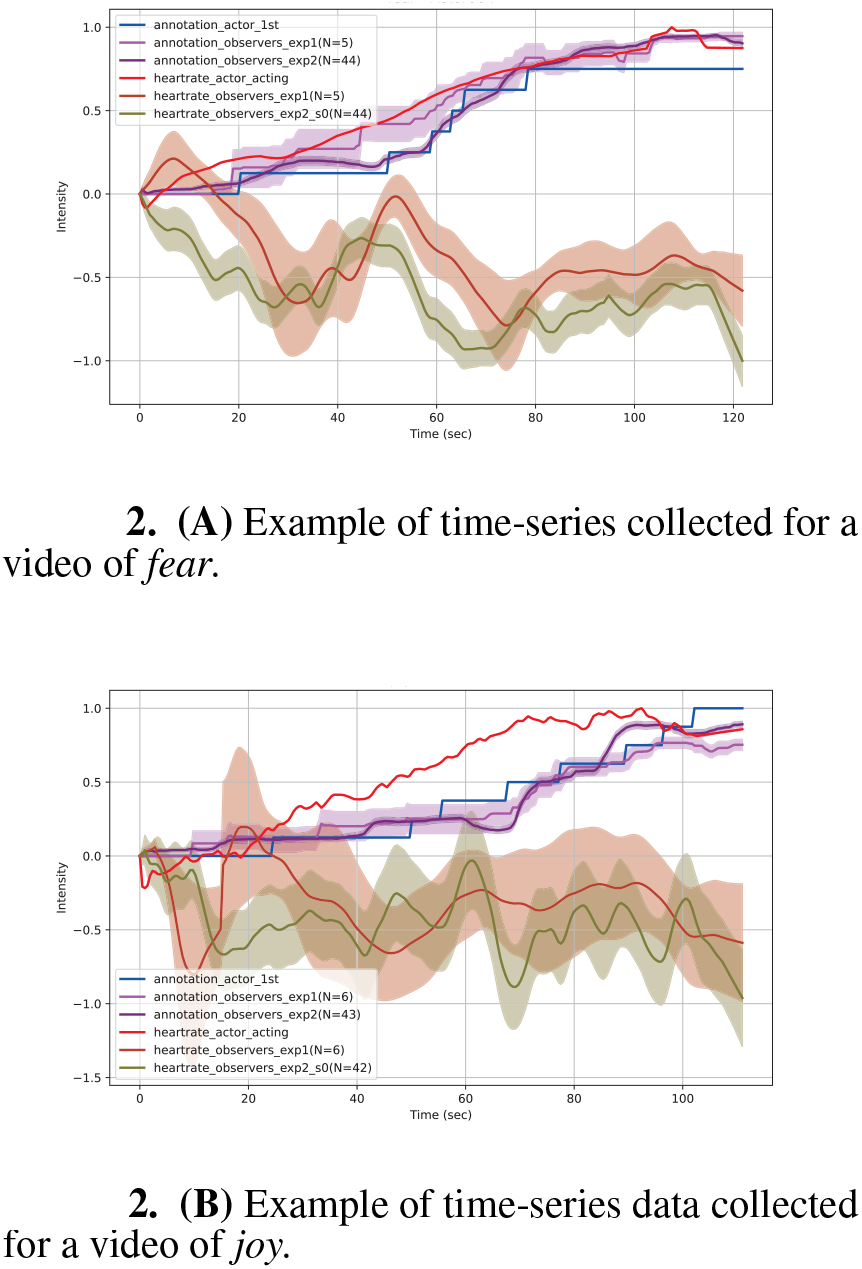
Example time-series data of the actor’s and observers’ annotations and heart rates for two videos: **(A) fear** and **(B) joy.** All heart rates have been shifted to start at a value of zero for clarity.

#### 3.4.2 Comparison of actors’ heart rate between acting and annotation phases, and between neutral and emotional (with and without) conditions

Figure 3 shows the mean HR across the three conditions during both acting and annotation. Table 2 presents the results of a linear mixed model (LMM), indicating that actors’ mean HR per video was higher during acting compared to annotation (*β* = 15.1, *t*(98) = 6.2, *p* < .001, CI = [10.4, 19.9]), and higher on with-expression condition compared to neutral condition (*β* = 4.11, *t*(98) = 2.06, *p* = .042, CI = [0.26, 7.97]). Elevated HR during acting was particularly evident for the with-expression condition (*β* = 7.27, *t*(98) = 2.84, *p* = .012, CI = [1.80, 12.74]).

**Table 2.** Results of linear mixed effect model on actors’ mean heart rate per video.

|  | **Estimate** | **Std. error** | **Df** | **t-value** | **p-value** | **2.5%** | **97.5%** |
| --- | --- | --- | --- | --- | --- | --- | --- |
| (Intercept) | 82.18 | 6.64 | 7.97 | 12.37 | < .001 | 68.52 | 95.83 |
| Acting | 15.13 | 2.45 | 98.00 | 6.18 | < .001 | 10.40 | 19.85 |
| With-expression | 4.11 | 2.00 | 98.00 | 2.06 | .042 | 0.26 | 7.97 |
| Without-expression | 2.46 | 2.00 | 98.00 | 1.23 | .22 | -1.39 | 6.32 |
| With-expression:Acting | 7.27 | 2.84 | 98.00 | 2.57 | .012 | 1.80 | 12.74 |
| Without-Expression:Acting | 3.25 | 2.83 | 98.00 | 1.15 | .25 | -2.21 | 8.70 |

**Table 3.** Results of task-wise linear mixed effect model on actors’ mean heart rate.

| (a) Acting phase |  |  |  |  |  |  |  |
| --- | --- | --- | --- | --- | --- | --- | --- |
|  | **Estimate** | **Std. error** | **Df** | **t-value** | **p-value** | **2.5%** | **97.5%** |
| (Intercept) | 97.31 | 7.13 | 7.82 | 13.65 | < .001 | 82.63 | 111.98 |
| With-expression | 11.27 | 2.09 | 45.00 | 5.40 | < .001 | 7.18 | 15.36 |
| Without-expression | 5.71 | 2.08 | 45.00 | 2.75 | .0085 | 1.65 | 9.77 |
| (b) Annotation phase |  |  |  |  |  |  |  |
|  | **Estimate** | **Std. error** | **Df** | **t-value** | **p-value** | **2.5%** | **97.5%** |
| (Intercept) | 82.18 | 6.15 | 7.18 | 13.36 | < .001 | 69.43 | 94.93 |
| With-expression | 4.11 | 0.85 | 46.00 | 4.82 | < .001 | 2.44 | 5.78 |
| Without-expression | 2.46 | 0.85 | 46.00 | 2.89 | .0059 | 0.79 | 4.13 |

**Figure 3.**
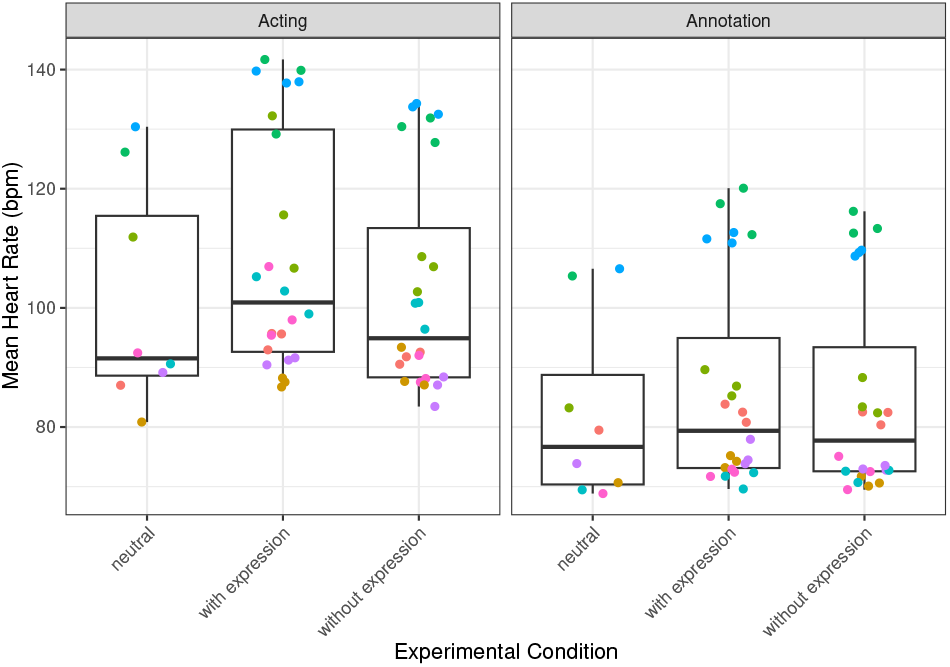
Mean heart rate (HR) of *actors* during acting (left) and annotation (right) for neutral episodes (left), emotional episodes with overt expressions (middle), and emotional episodes without overt expressions (right). Each point represents the average HR for one video, and colors differentiate individual actors.

To further examine the main effects within each phase, separate LMMs were performed for acting and annotation. Although the without-expression condition did not differ significantly from neutral overall, the phase-wise results (Table 2) showed higher mean HR in both emotional conditions compared to neutral: (a) in the acting phase, for the with-expression condition (*β* = 11.3, *t*(45) = 5.4, *p* < .001, CI = [7.2, 15.4]) and for the without-expression condition (*β* = 5.7, *t*(45) = 2.8, *p* = .009, CI = [1.6, 9.8]); (b) in the annotation phase, for the with-expression condition (*β* = 4.1, *t*(46) = 4.8, *p* < .001, CI = [2.4, 5.8]) and for the without-expression condition (*β* = 2.5, *t*(46) = 2.9, *p* = .006, CI = [0.8, 4.1]).

#### 3.4.3 Temporal changes in actors’ heart rate between first and second halves

Figure 4 presents the mean heart rate (HR) for the first and second halves of the videos. Shapiro–Wilk normality tests indicated that HR distributions for both halves in both phases deviated from normality: acting phase—first half (*W* = 0.880, *p* = .0085), second half (*W* = 0.831, *p* < .001); annotation phase—first half (*W* = 0.790, *p* < .001), second half (*W* = 0.796, *p* < .001). Therefore, Wilcoxon signed-rank tests were used.

**Figure 4.**
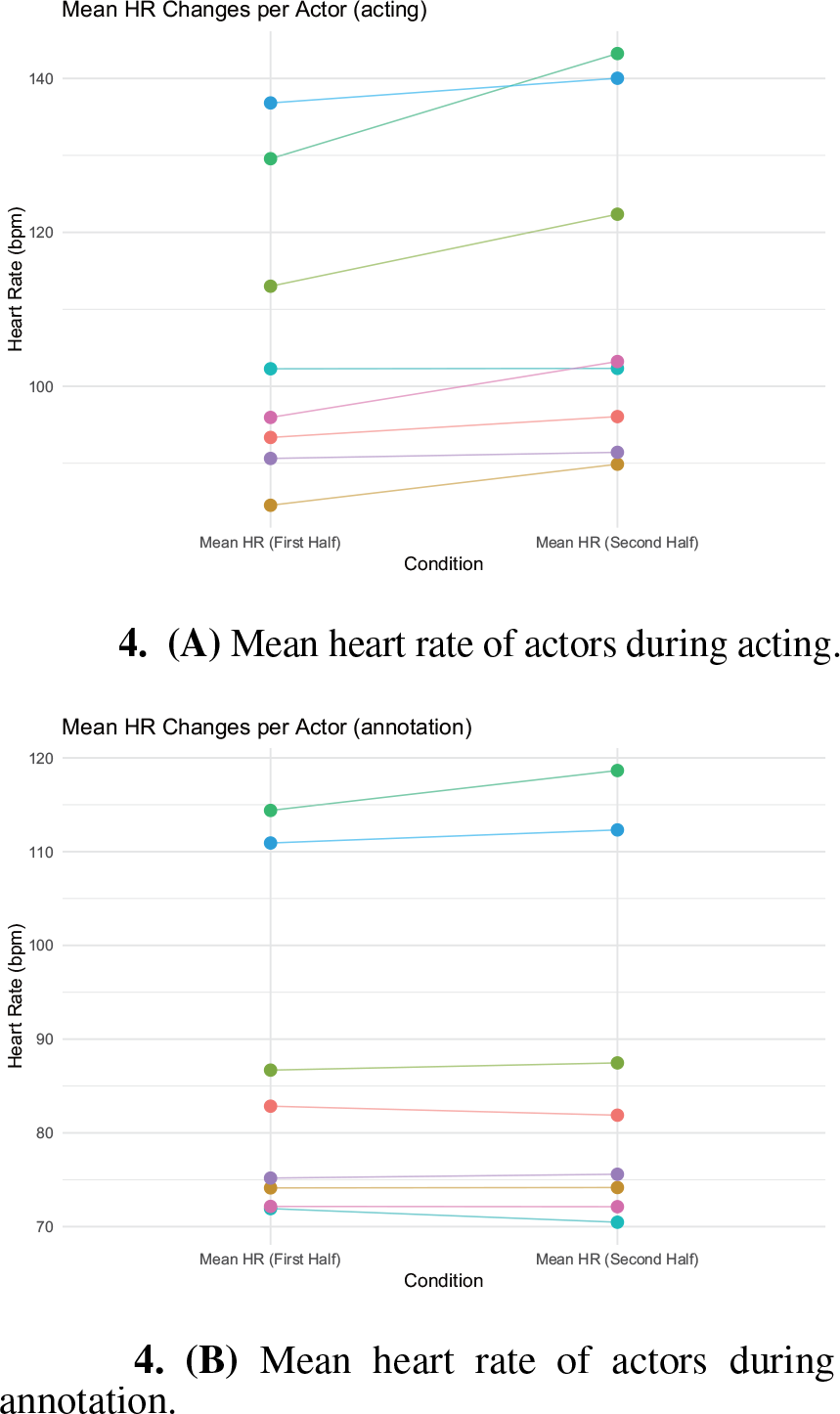
Mean heart rate per *actor* for the first and second halves of each video while **(A) acting** and **(B) rating.** Each dot represents heart rate of an individual actor for a single video, while colors indicate actors. An upward slope indicates an increase in heart rate, while a downward slope indicates a decrease. This visualization highlights the direction and magnitude of heart rate changes across actors.

During the acting phase, actors’ HR increased from the first half (mean = 105.8, median = 100.7) to the second half (mean = 111.1, median = 102.5) across the 24 emotional-expression videos (*V* = 9, *p* < .001, *r* = .822), indicating a marked rise in physiological arousal as actors progressed through their performances. In contrast, during the annotation phase, no significant difference was observed between the first half (mean = 86.0, median = 79.8) and the second half (mean = 86.6, median = 79.0) (*V* = 125, *p* = .484, *r* = .146), suggesting that HR remained stable and that physiological engagement was lower than in the acting phase.

For completeness, per-emotion summaries of within-video HR change (computed as the percentage change from the within-video minimum to maximum HR) showed the smallest median increase for *surprise* (12.7%), followed by *disgust* (15.1%), *joy* (18.3%), *fear* (20.1%), *anger* (22.6%), and the largest for *sadness* (28.5%).

### 3.5 Discussion

Taken together, the temporal analyses indicate that overt acting is accompanied by progressive HR increases within videos, whereas HR remains comparatively stable during annotation. Although HR is an imperfect stand-alone proxy for felt emotion, the common assumption that HR indexes arousal and the frequently observed V-shaped valence–arousal relationship (Kuppens et al., 2013) support interpreting these increases as successful elicitation of the intended affect during performance.

Our emotion-wise ordering—smallest HR change for *surprise* and largest for *sadness*—is consistent with adjacent literatures methodologically close to acted affect. First, studies using directed facial action and intentional affect generation, conducted with both professional actors and non-actor samples, have reported greater cardiac activation for anger and fear—and in some protocols, sadness—than for disgust or surprise (Levenson et al., 1990). Second, the comparatively larger HR changes we observed for sadness plausibly reflect bereavement-themed narratives in our corpus, consistent with evidence that grief-related states elevate HR and reduce HRV (O’Connor et al., 2002). Third, actor-focused resources indicate that physiology recorded during enacted narratives tracks arousal in ways useful for affect analysis (Aly et al., 2024). This directionality is further supported by review evidence showing that emotion-differentiated autonomic patterns depend on elicitation method and time course, yet typically yield greater cardiac activation for anger and fear than for disgust or surprise (Kreibig, 2010a).

Interestingly, for videos without emotional expression, the difference between acting and annotation phases was smaller than for videos with expression, suggesting that suppressing expressivity may attenuate physiological arousal relative to overt expression. Although collecting actor physiology during self-report, in addition to during acting, may further illuminate these effects, that inquiry lies beyond the scope of this paper.

## 4 OBSERVER EXPERIMENT 1: A LITERATURE FOLLOW-UP

In this section, the autobiographical videos with expressed emotions created in Section 3 were evaluated by external observers. Observers completed an empathic accuracy task in which they provided time-continuous ratings of the target’s emotional intensity while viewing the videos; these annotations were then analyzed alongside individual IRI scores. Following established empathic accuracy protocols (Mackes et al., 2018; McKenzie et al., 2022), we (i) quantified empathic accuracy (EA) from the time-continuous ratings and (ii) examined associations between EA and observers’ dispositional empathy (IRI subscales). Our aim was to verify that the newly created video stimuli reproduce the behavioral patterns reported in prior work (Mackes et al., 2018), thereby establishing comparability with the existing literature.

However, as in prior studies and in our replication (Experiment 1), observers annotated emotional intensity without being informed of the video’s target emotion category. The absence of a predefined category limits interpretability because the specific emotion referenced by a given intensity trace remains ambiguous. To address this limitation, Experiment 2 adopted a revised protocol in which the target category was disclosed prior to viewing to guide the annotation process.

### 4.1 Participants (observers)

Twelve female college students in their 20s were recruited; all reported no current psychiatric treatment needs and were judged to be psychologically stable, given that some stimuli could potentially trigger past traumas. Recruitment occurred from February 24 to March 16, 2023. On the day of data collection, the experimenter read aloud a written information sheet and consent form, answered any questions, and obtained written informed consent from all participants before the experimental procedure began.

### 4.2 Procedure

Each participant completed three tasks: (i) a personality questionnaire, (ii) a time-continuous emotion-intensity annotation task with concurrent physiological recording, and (iii) post-annotation questions for each assigned video.

#### 4.2.1 Personality questionnaire

As with the actors, observers completed the IRI-J via an online form prior to the experimental session.

#### 4.2.2 Time-continuous emotion-intensity annotation

Each observer viewed a randomly assigned set of 14 videos (from the 24 autobiographical clips featuring emotional expression), once each, in random order, with a one-minute break after the seventh video. While watching each video for the first and only time, observers continuously rated the intensity of the emotion portrayed using a 9-point scale (anchored at 0 = no emotion) via a mouse.

Physiological signals were recorded concurrently with annotation, mirroring the actors’ recording setup. ECG and EDA were measured with the Intercross-415 system (Intercross Corp.), and three-channel frontal EEG was recorded. A five-minute, eyes-open resting baseline preceded the annotation block. EDA and EEG were collected for completeness but were not analyzed in the present study.

#### 4.2.3 Post-annotation questions per video

After each video, participants answered three questions: (Q1) their own current emotional state (chosen from neutral, joy, sadness, disgust, anger, fear, and surprise); (Q2) the extent to which they felt the same emotion as the person in the video (1 = not at all to 9 = very much so); and (Q3) the ease of evaluating the video’s emotional intensity (1 = very difficult to 9 = very easy). Note that none of these responses contributed to the EA score, and Q2 was answered without being informed of the target emotion category. Q1 follows the question used in the literature(Mackes et al., 2018). Q2 and Q3 were included to assess perceived emotional clarity and appraisal difficulty, but these items were not analyzed in the present study.

### 4.3 Analysis

Following the analysis of previous studies (Mackes et al., 2018; McKenzie et al., 2022), Pearson’s correlation coefficient *r* was calculated between the emotional intensity annotations provided by actors and those by observers to derive the EA scores. Cases where the variance in emotional intensity evaluation was zero, rendering the calculation of EA scores unfeasible, were excluded from the analysis. Additionally, data with EA scores exceeding *±*2 standard deviations from the group mean were also removed to ensure robust analysis. This criteria excluded one participant; she had a history of arrhythmia. Subsequently, the EA scores were transformed from the original coefficient *r* to Fisher’s Z (Fisher, 1915) (Z-EA scores), to make results comparable to those reported as Z-EA scores in the literature (Mackes et al., 2018). Further, an analysis of variance was conducted on the Z-EA scores to explore differences across emotional categories.

We then examined whether the Z-EA scores differed between cases where observers’ reported emotion matched the actor’s instructed category and those that did not in Q1.

Consistent with prior analyses (Mackes et al., 2018), we correlated each participant’s mean Z-EA score with the IRI subscales.

To test whether actors’ physiological arousal predicted empathic accuracy, we next fit a linear mixed-effects model with Z-EA as the dependent variable. Fixed effects were smoothed heart-rate rate-of-change, emotion category, and their interaction; Observer ID was included as a random intercept. Emotion category was dummy-coded with *surprise* (the condition showing the strongest positive slope) as the reference level.

Finally, analogous to the actors’ heart-rate analysis, we compared observers’ mean heart rate during annotation between the first and second halves of each video to assess overall trends in emotional arousal across observers.

### 4.4 Results

#### 4.4.1 Qualitative evaluation of observers’ annotations and heart rate

Figure 2 illustrates observers’ HR data and time-continuous emotion intensity annotations from Experiment 1 (during video viewing). Most observers’ annotations broadly tracked both the actors’ HR dynamics and the actors’ self-annotations, whereas observers’ average HR showed no consistent pattern and exhibited substantial inter-individual variability.

#### 4.4.2 Empathic accuracy scores: (*Z*-)EA scores overall and by emotion category

The overall empathic accuracy (EA) matched previously reported high levels: the mean correlation was *r* = .77 and the mean within-individual standard deviation was *iSD* = .23, compared to *r* = .75 with *iSD* = .35 (Mackes et al., 2018) and *r* = .81–.84 (McKenzie et al., 2022). Note that for (McKenzie et al., 2022), *r* values were back-transformed from Fisher’s *Z*-transformed EA scores, and *iSD* was not reported.

A Wilcoxon rank-sum test indicated that Z-EA scores for *real* pairs (actor–observer annotations from the same video) were higher than for *pseudo* pairs (randomly matched actor–observer annotations from different videos with length differences *≤* 5 seconds): real, 1.26 *±* 0.34 (*SD*; *n* = 129) vs. pseudo, 1.06 *±* 0.23 (*n* = 138); *W* = 11,170, *p* < .001.

A repeated-measures ANOVA on *Z*-EA scores revealed a main effect of emotion category, *F* (5, 45) = 5.681, *p* < .001, *η*^2^ = .152. Mean *Z*-EA by category was as follows: *sadness* 1.53 *±*0.35, *joy* 1.41*±* 0.40, *anger* 1.39 *±*0.42, *fear* 1.37 *±*0.65, *disgust* 0.96 *±*0.61, and *surprise* 0.91 *±*0.58. Shaffer-adjusted post hoc comparisons indicated that *Z*-EA for *surprise* was lower than for *sadness, joy*, and *anger* (all *p* < .046). Briefly situating these results in prior EAT work: *sadness* yielded the highest *Z*-EA here and in McKenzie et al. (2022), with *joy*/*happiness* and *fear* in an intermediate range in both; by contrast, the ordering for *anger* differs across studies (lower in McKenzie et al. (2022), especially for ASD), and Mackes et al. (2018) contrasted only *happy* vs. *sad*—reporting higher *Z*-EA for *happy* than *sad*. We return to possible reasons for these divergences in 4.5.

#### 4.4.3 *Z*-EA for observer–actor emotional sharing versus non-sharing

As a comparative validation following prior work that used observers’ Q1 responses (Mackes et al., 2018), we computed the proportion of trials in which the observer’s reported emotion matched the actor’s instructed category. Across all trials, observers matched the actor’s category in 56.0% of cases; agreement was highest for *joy* (68.0%) and lowest for *anger* (34.0%). A Wilcoxon rank-sum test revealed no significant difference in *Z*-EA between trials where the observer’s Q1 matched the actor’s instructed category (1.30 *±*0.27, *n* = 77) and trials where it did not (1.22 *±*0.40, *n* = 58); *W* = 2,377, *p* = .52, consistent with prior research (Mackes et al., 2018).

#### 4.4.4 Relationship between *Z*-EA and observers’ personality traits

Consistent with previous research (Mackes et al., 2018), participants’ mean *Z*-EA scores correlated positively with Perspective Taking (PT) on the IRI-J (*r* = .54, *p* < .05). In contrast to prior findings, mean *Z*-EA correlated negatively with Personal Distress (PD), *r*(8) = *−*.71, *p* < .05. When analyses were restricted to negative-emotion videos (anger, disgust, fear, sadness), these associations were stronger (PD: *r*(8) = *−*.77, *p* < .01; PT: *r*(8) = .67, *p* < .05). No other IRI subscales were significantly related to mean *Z*-EA (all *p* > .60).

#### 4.4.5 Relationship between *Z*-EA and actors’ heart rate change during acting

Emotion-wise mean *Z*-EA scores were positively correlated with actors’ within-video heart-rate (HR) change during acting (maximum minus minimum; see Section 3.4.3), *r*(4) = .880, *p* = .021. To examine category-specific effects, a linear mixed-effects model revealed a significant interaction between HR change and emotion category (*p* < .01). Simple-slope estimates showed that increases in actors’ HR predicted higher empathic accuracy (*Z*-EA) in *surprise* (reference; *β* = 3.47, *p* < .001) and in the other categories except *anger* (range *β* = 0.56–1.14; all *p* < .05). In contrast, for *anger*, greater physiological arousal predicted lower empathic accuracy (*β* = *−*0.45, *p* < .01). Notably, this divergence for *anger* parallels cross-study differences in EAT performance patterns and is revisited in the Discussion alongside comparisons to Mackes et al. (2018); McKenzie et al. (2022).

#### 4.4.6 Observers’ heart-rate change during annotation

Figure 5 depicts the change in observers’ heart rate from the first to the second half of each video. In contrast to the actors’ pattern, most observers showed lower HR in the second half, consistent with the inferential results. Shapiro–Wilk tests indicated departures from normality for both halves (first half: *W* = 0.972, *p* = .0023; second half: *W* = 0.946, *p* < .001). Accordingly, a Wilcoxon signed-rank test revealed a significant decrease in HR from the first half (mean = 78.1, median = 78.3) to the second half (mean = 77.0, median = 77.1), *V* = 5,121, *p* = .0035, *r* = .249 (small–to–moderate effect).

**Figure 5.**
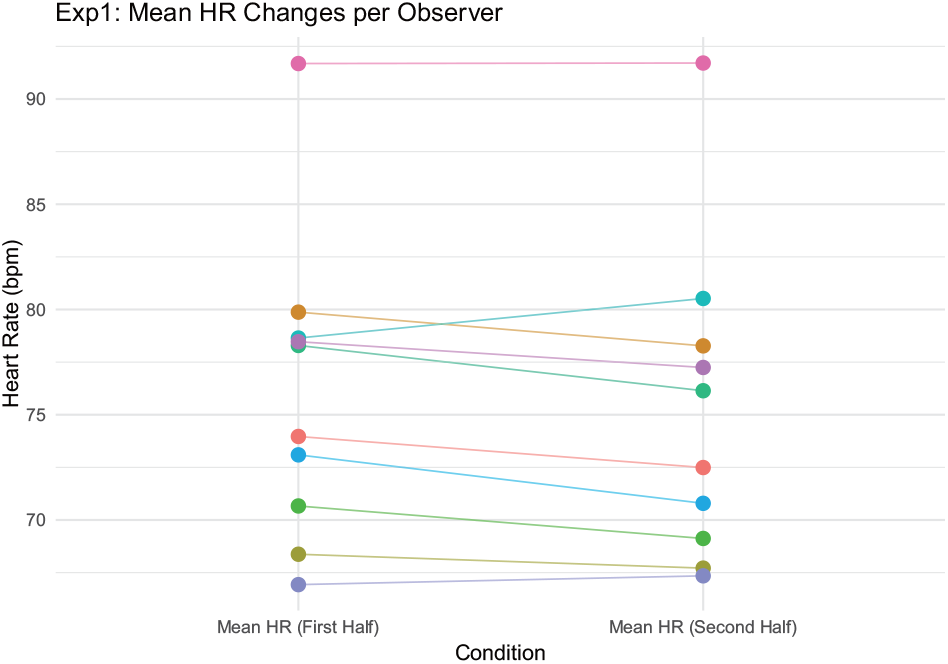
Mean heart rate per *observer* for the first and second halves while rating to videos with emotional expressions in *Experiment 1*. Colors indicate observers.

**Figure 6.**
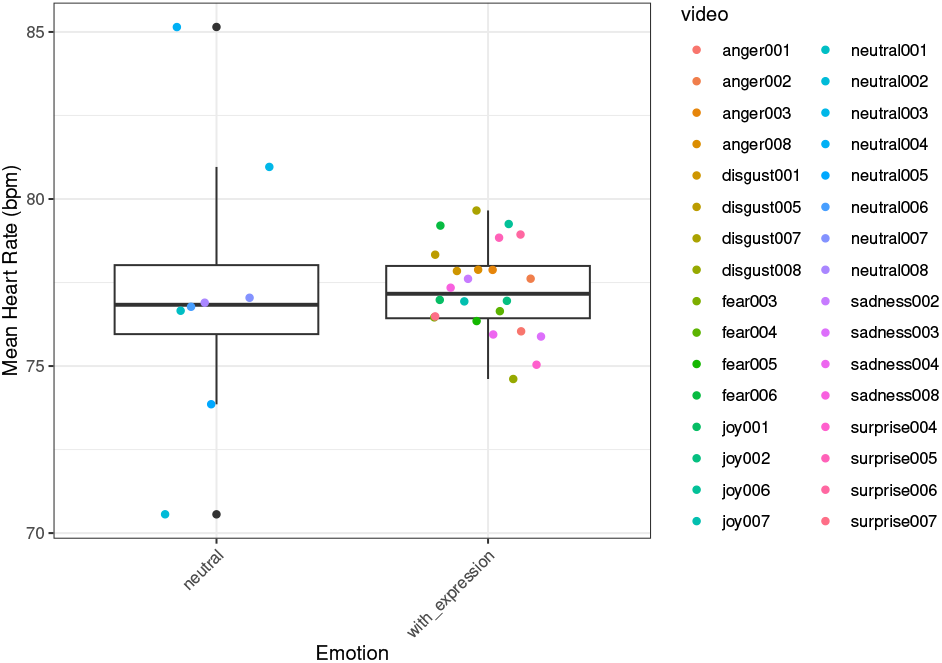
Mean HR of *observers* during for neutral episodes (left), emotional episodes with emotional expressions (right) in *Experiment 2*. Plots are mean per video.

### 4.5 Discussion

#### 4.5.1 Validity of empathic accuracy scores in comparison with the literature

Overall empathic accuracy (EA) in Experiment 1 fell between prior reports: mean *r ≈*.77 vs. *r* = .75 in Mackes et al. (2018) and *r* = .81–.84 in McKenzie et al. (2022), indicating that the present stimuli were suitable for empathic-accuracy judgments. Emotion-wise analyses further showed lower *Z*-EA for *surprise* than for *anger, fear, joy*, and *sadness*.

Situating our category pattern relative to prior EAT reports, McKenzie et al. (2022) likewise found the highest *Z*-EA for *sadness*, with *happiness* and *fear* in an intermediate range, whereas *anger* was comparatively lower at least in their ASD group. By contrast, Mackes et al. (2018) contrasted only *happy* vs. *sad* and reported higher accuracy for *happy*. Thus, our rank order (top *sadness*; *joy*/*fear* intermediate) aligns with McKenzie et al. (2022) but departs from the two-emotion contrast in Mackes et al. (2018). We return to possible methodological and stimulus-level reasons for these differences below.

As a bridge to those explanations, we note that the emotion labels themselves may not be the sole drivers of category means. In our data, an index of actors’ autonomic activation during acting covaried with *Z*-EA across categories, pointing to arousal-linked accounts rather than purely discrete-emotion accounts.

Crucially, the emotion-wise mean *Z*-EA was positively associated with actors’ within-video heart-rate (HR) change during acting (second minus first half; see Section 3.4.3), *r*(4) = .880, *p* = .021. Given longstanding evidence that HR indexes autonomic arousal (Kreibig, 2010a) and the typical V-shaped valence–arousal relation (Kuppens et al., 2013), this pattern suggests that between-video differences in *enacted arousal*—rather than the discrete emotion label per se—contributed to the observed category differences in *Z*-EA. In other words, videos that elicited larger autonomic responses in the actors tended to yield higher empathic accuracy, which is theoretically coherent because observers were instructed to track *intensity*. The relatively low *Z*-EA for *surprise* is therefore plausibly attributable to its lower sustained arousal (and more transient time course) in our stimuli, rather than an intrinsic deficit in reading “surprise.”

We note that this correlation does not establish causality and that HR is an imperfect proxy for arousal; moreover, arousal and categorical content were not independently manipulated here. Nevertheless, the convergence between actors’ physiological responses and observers’ intensity-tracking accuracy supports the validity of these stimuli for EA research. A fruitful next step would be to test a “mediation-by-arousal” account more directly—for example, by predicting trial-level *Z*-EA with a mixed-effects model that includes per-video HR change as a covariate (alongside emotion category, video length, actor ID, and observer traits), or by experimentally varying arousal independently of category.

#### 4.5.2 Relationship between (*Z*-)EA and observers’ personality traits

Prior work has linked EA more strongly to cognitive than affective components of empathy (Mackes et al., 2018), based on both neural activation during annotation and correlations with the IRI Perspective Taking (PT) scale. We replicated this association: mean *Z*-EA correlated positively with PT. In addition, mean *Z*-EA correlated negatively with Personal Distress (PD), an effect that was particularly strong for negative-emotion videos. This suggests that a tendency to experience self-oriented distress in response to others’ suffering may hinder precise moment-to-moment tracking of another’s emotional intensity, consistent with the view that empathic accuracy relies more on cognitive perspective-taking than on affective resonance.

Consistent with previous studies, we also observed no difference in *Z*-EA between trials where observers reported the same emotion category as the actor and those where they did not, indicating that sharing the same labeled emotion is not necessary for (and may not enhance) intensity-based empathic accuracy. These findings support the validity of the EMPAC stimuli for EA research, while also highlighting a limitation of the protocol: it does not measure accuracy with respect to intended emotion *categories* and provides only an indirect assessment of cognitive empathy.

#### 4.5.3 Emotion-specific reversal for anger in the relationship between *Z*-EA and actors’ heart-rate change

Although greater physiological arousal in actors was generally associated with higher empathic accuracy across categories, this trend was *reversed* for *anger*: greater HR change predicted *lower Z*-EA. One plausible explanation lies on the observer side: anger can reduce perspective-taking and increase heuristic, top–down processing (Yip and Schweitzer, 2019; Lerner and Tiedens, 2006), potentially disrupting the fine-grained tracking required for *Z*-EA. On the target side, HR is a coarse index of arousal and lacks specificity for anger (Kreibig, 2010a). Thus, increased HR in angry episodes may not offer clear, diagnostic cues for observers to match, leading to reduced empathic accuracy in this condition. Importantly, this anger-specific divergence resonates with cross-study differences in EAT performance patterns—for instance, comparatively lower anger accuracy in parts of McKenzie et al. (2022)—and helps contextualize why our category ordering aligns with McKenzie et al. (2022) for *sadness*/*fear*/*joy* but diverges for *anger*.

#### 4.5.4 Inverse HR-change trend between actors and observers

Among observers, HR decreased from the first to the second half of videos, in contrast to the actors’ increase. One functional interpretation is that HR deceleration indexed sustained attention and task engagement during moment-to-moment intensity tracking (Lacey and Lacey, 1974; Jennings and Coles, 1991). Crucially, this pattern need not imply reduced engagement; rather, it may reflect an attentionally tuned state that facilitates precise annotation.

This interpretation yields testable predictions. First, at the trial level, greater HR deceleration during annotation should be associated with higher *Z*-EA after controlling for video-, actor-, and observer-level random effects (e.g., via a mixed-effects model using the within-trial HR slope or second–minus–first-half change as a covariate). Second, experimentally manipulating attentional load should causally affect both physiology and accuracy: adding a light secondary task (e.g., 1-back tones) should reduce *Z*-EA and systematically alter cardiac dynamics during annotation if attention is a limiting resource. Third, separating sympathetic and parasympathetic contributions would sharpen interpretation: concurrent indices such as high-frequency HRV/RSA (parasympathetic), pre-ejection period (sympathetic), EDA (arousal), and pupil diameter (cognitive load) can disambiguate whether the observed HR decreases primarily reflect vagal engagement rather than low arousal. Finally, orthogonally manipulating actor arousal (e.g., low/medium/high intensity within the same category) would permit a more direct test of a mediation pathway in which variation in actor arousal modulates observer attention (indexed by HR deceleration/RSA), which in turn affects empathic accuracy.

Together with the actor-side finding that videos eliciting larger autonomic responses tended to yield higher *Z*-EA, these observer-side proposals outline a principled path to adjudicate whether attention-indexed cardiac deceleration is merely epiphenomenal or functionally supports empathic intensity tracking.

## 5 OBSERVER EXPERIMENT 2: CATEGORY-SPECIFIC EMPATHIC ACCURACY

In Experiment 2, we re-evaluated empathic accuracy for the autobiographical videos with emotional expression from Experiment 1 using a revised annotation procedure. The key change was that, whereas in Experiment 1 observers were not told the target emotion, in Experiment 2 they were informed in advance of the actor-instructed emotion category and asked to provide time-continuous intensity ratings for that specified target. Importantly, as noted in Experiment 1, we treated the actors’ annotations as category-specific because each clip was *assumed* to express a single, discrete emotion in accordance with the acting instructions, while acknowledging that some narratives may incidentally contain mixed affect. Accordingly, the same actor annotations served as the target time series in both experiments without modification.

Finally, given the modest observer sample in Experiment 1 (mean *≈*5.6 observers per video) and clear precedents in prior work (Mackes et al., 2018; McKenzie et al., 2022), Experiment 2 increased the number of participants to strengthen category-specific analyses.

### 5.1 Participants (observers)

Using the same inclusion criteria as in Experiment 1, we recruited 87 healthy university students (44 women, 21.0 *±*1.6 years; 43 men, 20.9 *±*2.1 years; overall age 21.0 *±*1.9 years). After data cleaning (excluding trials with missing heart-rate data and their associated annotations), the final sample yielded an average of 42.9 *±* 0.95 annotators per emotional video (range: 41–45), substantially larger than in Experiment 1 (mean *≈* 5.6 per video).

Recruitment occurred from December 6 to December 13, 2023. On the day of data collection, the experimenter read aloud a written information sheet and consent form, answered any questions, and obtained written informed consent from all participants prior to the experimental procedure.

### 5.2 Procedure

Each participant completed four tasks: (i) a practice session for time-continuous annotation, (ii) a time-continuous emotion-intensity annotation task with concurrent physiological recording, (iii) an overall emotion-judgment task for each assigned video, and (iv) personality questionnaires.

#### 5.2.1 Practice session (continuous-annotation calibration)

To accommodate individual differences in time-continuous annotation, participants first familiarized themselves with the interface and then completed a perceptual tracking task to characterize their slider response tendencies (e.g., gain, lag). Following previous methods (Booth and Narayanan, 2020), participants viewed a video with continuously varying green luminance and tracked its brightness using a vertical slider (range: 0 to 1; 0 = lowest brightness, 1 = highest). Two training videos were used: one with linear luminance change and one with a sigmoidal trajectory. Practice data were not analyzed further.

#### 5.2.2 Time-continuous emotion-intensity annotation

Each participant viewed a pseudo-random set of 13 videos in randomized order: 12 emotional clips from the 24 autobiographical, overt-expression videos (6 emotions *×*4 actors) and 1 neutral clip (from the eight actors). Before each video, the target emotion category (actor-instructed) was disclosed, and participants continuously rated the intensity of that specified emotion while viewing, using the same slider (0 to 1; 0 = no emotion). Each video was presented twice consecutively with a one-minute pause before replay, and a 10-minute break followed the sixth video. This design ensured uniform exposure and supported reliable estimation of empathic accuracy across emotion categories.

#### 5.2.3 Overall emotion judgment per video

After the initial viewing of each video, participants answered ten questions. The first five referred to *their own* state and the next five to the *target actor’s* state: (1) emotion category (neutral, joy, sadness, disgust, anger, fear, surprise), (2) confidence in the response to Question 1, (3) valence, (4) arousal, and (5) dominance. Questions 2–5 were answered on a visual analogue scale. After completing these items, the intended emotion category of the video (i.e., the actor-instructed category) was revealed. Participants then viewed the video a second time and continuously annotated *that target emotion* while watching.

#### 5.2.4 Physiological measurement

ECG and EDA were recorded throughout the viewing sessions using a Polymate Pocket MP208 (Miyuki Giken).

#### 5.2.5 Personality questionnaires

As in Experiment 1, observers completed the IRI-J via an online form. They also completed the Japanese versions of the Ten Item Personality Inventory (TIPI-J) (Oshio et al., 2012) and the Autism-Spectrum Quotient (AQ-J) (Wakabayashi et al., 2004). To focus on comparisons between Experiments 1 and 2, only IRI scores were analyzed; TIPI and AQ data were collected but not analyzed in the present study.

### 5.3 Analysis

To further validate the actors’ emotional portrayals from a complementary perspective, we first computed an accuracy metric—defined as the proportion of correct responses, i.e., trials in which an observer’s categorical judgment (Question 1 for the target actor) matched the actor-instructed emotion. Accuracy was calculated at two levels: (i) the individual-observer level and (ii) the video level based on the majority vote across observers. Next, we directly compared participants’ annotations in Experiments 1 and 2, then calculated their EA scores and contrasted them with scores from Experiment 1. We also examined whether the observers’ EA scores correlated with their personality traits (IRI scores), replicating the approach used in Experiment 1. Furthermore, we investigated any potential gender effects on the observers’ EA scores, which was not applicable to Experiment 1, which used female participants only. Finally, mirroring the procedure in Experiment 1, we compared average observer heart rates between the first and second halves of the videos to determine whether the videos influenced observers’ heart rates.

Finally, as in Experiment 1, we tested whether actors’ HR changes predicted observers’ *Z*-EA. We fit a linear mixed-effects model with fixed effects for smoothed HR change, emotion category, and their interaction, and observer-specific random intercepts. Emotion category was dummy-coded with *surprise* as the reference level.

### 5.4 Results

#### 5.4.1 Qualitative evaluation of observers’ annotations and heart rate

Figure 2 illustrates observers’ HR data and time-continuous intensity annotations from Experiment 2 (during video viewing). As in Experiment 1, most observers’ annotations broadly tracked both the actors’ HR dynamics and the actors’ self-annotations. Although observers’ average HR showed no consistent pattern and exhibited considerable inter-individual variability, the overall HR patterns resembled those in Experiment 1.

#### 5.4.2 Observers’ accuracy in overall categorical emotion judgment

Observers’ categorical judgments (Question 1 for target actors) were correct on 86% of trials at the individual-observer level. At the video level, the majority label across observers matched the actor-instructed emotion in 100% of cases. These findings support the conclusion that actors’ intended emotions were effectively conveyed.

#### 5.4.3 Empathic accuracy scores: (*Z*-)EA scores overall and by emotion category

Mean observer annotations for each emotional video in Experiment 2 (*N* = 48.7 per video on average) were highly similar to those in Experiment 1 (*N* = 5.6), Pearson’s *r* = 0.95 *±* 0.03. However, the overall individual-level EA score in Experiment 2 was slightly higher than in Experiment 1 (mean *r* = .81, mean *iSD* = .19 *vs. r* = .77, *iSD* = .23), suggesting that disclosing the target emotion category reduced ambiguity for participants. Furthermore, the across-observer average annotation for each video showed an even higher correlation with the actors’ annotation (mean *r* = .86), reinforcing the validity of both observer and actor annotations.

A repeated-measures ANOVA on *Z*-EA scores revealed a main effect of emotion category, *F* (5, 83) = 28.57, *p* < .001, *η*^2^ = 0.256. Consistent with Experiment 1, emotion-wise mean *Z*-EA scores were positively associated with actors’ within-video HR change during acting (computed as the percentage change from the within-video minimum to maximum HR); see Section 3.4.3), *r*(4) = .757, *p* = .082. Mean *Z*-EA by category was as follows: sadness 1.55 *±* 0.23, joy 1.52*±* 0.10, anger 1.39*±* 0.17, fear 1.37 *±*0.36, disgust 1.02 *±* 0.33, and surprise 1.18 *±* 0.30. Shaffer-adjusted post hoc comparisons revealed that Z-EA scores for sadness and joy were significantly higher than those for disgust and surprise, while scores for anger and fear did not differ significantly.

#### 5.4.4 Relationship between *Z*-EA and observers’ personality traits

The observers’ mean Z-EA scores showed no correlation with their IRI subscales, differing from previous studies and from Experiment 1 (PD: *r*(85) = -.04, *p* = .72, EC: *r*(85) = .04, *p* = .72, PT: *r*(85) = .01, *p* = .89, FS: *r*(85) = .17, *p* = .12). We also found no gender effect on Z-EA scores, with *p >* .80 in a linear model Z-EA *∼* IRI_*s*_ + gender for all IRI subscales (indexed by *s*).

#### 5.4.5 Mean heart rate of observers (first versus second halves)

Figure 7 shows paired plots of observer heart rates in the first and second halves of the videos. Most participants demonstrated a decrease in HR from the first half (mean = 77.9, median = 77.9) to the second half (mean = 76.7, median = 76.4). A Wilcoxon signed-rank test yielded *V* = 386,043, *p* < .001, indicating a medium effect size (*r* = .373). Moreover, a Shapiro-Wilk normality test confirmed non-normal distributions for both halves (first half: *W* = 0.974, *p* < .001; second half: *W* = 0.974, *p* < .001), justifying use of the Wilcoxon test. The decreased observers’ HR suggests their cognitive engagement on the task, as discussed in Experiment 1.

**Figure 7.**
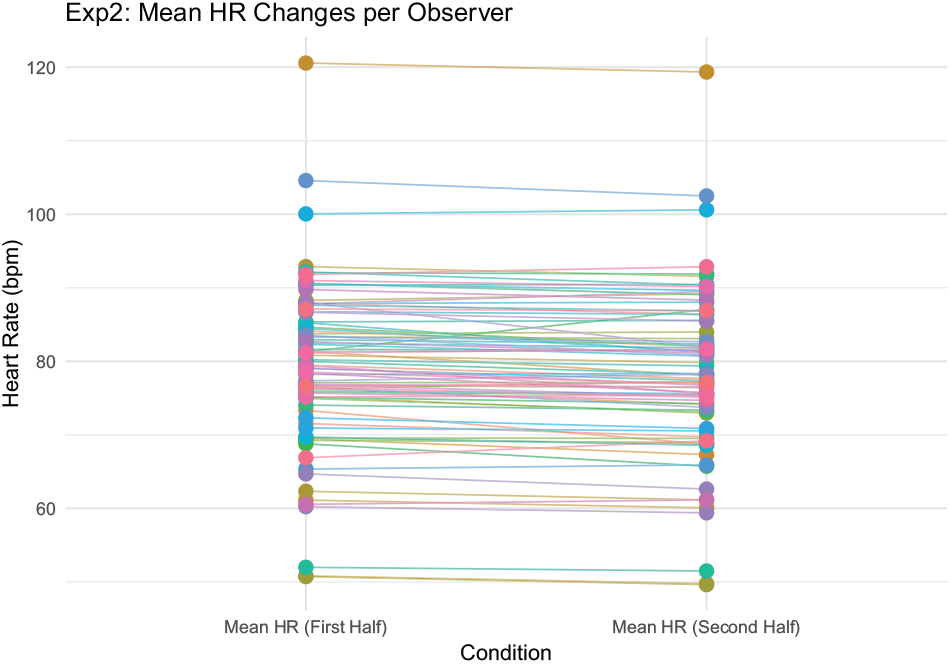
Mean heart rate per *observer* for the first and second halves while rating videos with emotional expressions in *Experiment 2*. Colors indicate observers.

#### 5.4.6 Relation between actors’ heart rate change and empathic accuracy

We applied the same model structure to Experiment 2 data. Again, we observed a significant interaction between heart rate change and emotion category (*ps <* .001). The strongest positive relationship was found in the *surprise* condition (*β* = 2.92, *p* < .001), while a negative association was again observed in the *anger* condition (*β* = *−*0.36, *p* < .001). All other emotions showed significantly positive slopes (*β*s = 0.33–1.21, all *ps <* .001).

### 5.5 Discussion

#### 5.1.1 Cross-experimental differences in annotation

Specifying the target category for time-continuous annotation yielded outcomes that differed from Experiment 1, which adhered to the protocol of McKenzie et al. (2022) and whose principal findings we replicated. Without knowledge of the correct category (Experiment 1), some observers misidentified the target emotion (overall categorical accuracy = 86%), which likely influenced their intensity annotations. At the same time, we note that additional procedural differences between Experiments 1 and 2 preclude perfectly direct comparisons.

#### 5.5.2 Relationship between Z-EA and actors’ arousal

A central, cross-experimental observation helps interpret these differences: the emotion-wise mean *Z*-EA scores were strongly associated with actors’ within-video HR change during acting (max–min), with *r* = .88 in Experiment 1 and *r* = .76 in Experiment 2. Given that HR is a conventional index of autonomic arousal (Kreibig, 2010a), these robust correlations suggest that the between-category variation in *Z*-EA primarily reflects differences in *enacted arousal* rather than discrete labels per se. In other words, videos in which actors exhibited larger cardiac responses tended to afford higher empathic accuracy in observers—coherent with the task instruction to track *intensity*. This account also explains why categories characterized by lower sustained arousal (e.g., *surprise*) showed comparatively lower *Z*-EA. We emphasize that these relationships are correlational and that HR is an imperfect proxy for arousal; nevertheless, their replication across experiments strengthens an arousal-based interpretation of the category effects.

Alternative explanations related to sample composition appear unlikely. Experiment 2 substantially increased the observer sample (mean *≈*48.7 observers per video vs. 5.6 in Experiment 1), yet Experiment 1 already replicated benchmark results from prior work employing larger samples (Mackes et al., 2018). Moreover, we found no effect of gender on *Z*-EA in Experiment 2, and gender differences were not reported in McKenzie et al. (2022). Thus, neither sample size nor gender distribution plausibly accounts for the differences between experiments.

A broader implication is that intensity judgments may rely on mechanisms that are partially distinct from those recruited for categorical (or dimensional valence-based) judgments. For example, amygdala circuitry is closely linked to perceived intensity, whereas orbitofrontal regions relate more to valence appraisal (Lewis et al., 2007); given the orbitofrontal cortex’s broad evaluative role (Fettes et al., 2017), these considerations underscore the value of methodological specificity and motivate examining empathy from multiple complementary perspectives. Our primary aim here is to introduce the dataset rather than arbitrate among annotation methods, but these preliminary patterns may inform future work.

Regarding observers’ physiology, as in Experiment 1 we observed a small decrease in HR from the first to the second half of videos, consistent with attention-related cardiac deceleration under sustained cognitive engagement (Lacey and Lacey, 1974; Jennings and Coles, 1991). Because observer-side autonomic changes are comparatively subtle and heterogeneous, they are harder to interpret than actors’ signals; nevertheless, including observer physiology is an important step toward a more complete account of affective empathy.

#### 5.5.3 Anger-specific reversal in the empathic accuracy–HR relationship

Across both experiments, greater target arousal (indexed by actors’ HR increase) generally predicted higher empathic accuracy, yet this association reliably reversed for *anger*: higher HR was linked to lower *Z*-EA, whereas other categories showed positive relationships. This anger-specific reversal is consistent with observer-side mechanisms whereby anger reduces perspective taking and promotes heuristic, top– down processing, disrupting the fine-grained, moment-to-moment attunement that *Z*-EA requires (Yip and Schweitzer, 2019; Lerner and Tiedens, 2006). A complementary target-side account is that HR is a coarse index of intensity/arousal with limited anger-specific diagnosticity due to physiological overlap across emotions (Kreibig, 2010b). Together, these mechanisms help explain why increases in actor HR can decouple observers’ ratings from targets’ rising intensity during anger, even as the same cue facilitates accuracy for other emotions.

## 6 GENERAL DISCUSSION

In this research, we investigated how *cognitive* and *affective* forms of empathy jointly shape observers’ understanding of another person’s emotions in a *time-continuous* paradigm. Across two experiments, we probed empathic accuracy (EA)—the alignment between a target’s self-reported emotional state and an observer’s inference—and analyzed whether subtle methodological changes (i.e., how observers annotate target emotions) might alter EA’s connection to observer traits such as *perspective-taking*. Our design was influenced by prior studies, including Mackes et al. (2018) and Jospe et al. (2020), both of which used naturalistic emotional stimuli and dynamic annotation paradigms, though each emphasized somewhat different routes (e.g., cognitive vs. affective empathy).

### 6.2 Replicating Mackes et al. in Experiment 1

Our *Experiment 1* closely mirrored the time-continuous emotion-intensity protocols introduced by Mackes et al. (2018). Observers annotated *broad emotional intensity* without being constrained to specific emotion labels. Like Mackes and colleagues, we found a *positive correlation* between EA scores and *perspective-taking*, suggesting that continuous tracking of someone’s fluctuating emotions primarily leverages *cognitive empathy*. This parallel underscores that subtle moment-to-moment evaluations can tap mentalizing mechanisms and provide robust empathic alignment—i.e., “reading” emotional curves in real time.

### 6.2 Changing Annotation Specificity in Experiment 2

In *Experiment 1*, observers provided *time-continuous emotion-intensity ratings* without any indication of which specific emotion the actor was portraying; they were free to judge the intensity of whatever emotion they perceived in the actor’s performance. By contrast, in *Experiment 2*, the *time-continuous annotation* remained in place, but observers were explicitly informed of the *emotion category* (e.g., anger or sadness) that the actor was attempting to convey. They then continuously rated the *perceived intensity* of that designated emotion over time. This subtle difference—shifting from an *open-ended* to a *category-defined* time-continuous rating—altered the relationship between empathic accuracy and observer traits such as perspective-taking. Specifically, while *Experiment 1* revealed a positive correlation between empathic accuracy and perspective-taking, as well as a strong negative correlation between empathic accuracy and personal distress, these associations were not observed in *Experiment 2*.

These findings suggest that *specifying* the target emotion may change the observer’s cognitive strategy, guiding them to verify or track that *one* assigned emotion rather than holistically inferring whichever emotion(s) they believe to be present. This result is consistent with arguments by Jospe et al. (2020), indicating that *annotation frames* (e.g., linguistic vs. visual cues, or open vs. category-defined rating instructions) can reallocate observers’ mental resources and modify which empathic routes they engage. When observers are constrained to focus on a single designated emotion, unstructured mentalizing and broader affective resonance may be reduced. This, in turn, could weaken the link between empathic accuracy and both perspective-taking and personal distress. However, these differences do not inherently favor one method over the other. Rather, the choice of annotation approach should align with the specific goals of a given study. The open-ended approach in *Experiment 1* may be more suitable for capturing spontaneous, holistic emotional interpretations, reflecting the observer’s natural empathic engagement. In contrast, the category-defined approach in *Experiment 2* may be better suited for studies aiming to assess accuracy in recognizing specific, pre-identified emotions — particularly in contexts where emotion recognition training or targeted empathy interventions are of interest. By highlighting how different annotation methods engage distinct cognitive and emotional processes, this study offers insight into selecting the most appropriate approach for future research. Researchers can tailor the annotation method to match their study’s objectives, whether prioritizing naturalistic emotional inference or focused accuracy on specific emotional categories.

### 6.3 Integrating Cognitive and Affective Empathy

Taken together, these experiments reinforce the notion that *cognitive* and *affective* components of empathy can be selectively emphasized by relatively minor changes in how observers rate or conceptualize the target’s emotion. Broad, *time-continuous* ratings (Experiment 1) facilitated higher empathic accuracy linked with perspective-taking, whereas *explicit* category-based ratings (Experiment 2) yielded different correlational patterns. Such findings confirm that “empathy” is not monolithic but rather arises through *varied interplay* between mentalizing (cognitive empathy) and sharing (affective empathy), depending on instructions and context.

### 6.4 Limitations

Despite these insights, several caveats should be noted:

1. **Stimulus Selection:** We used a limited number of autobiographical video clips, focusing on certain emotional events. Although this design is consistent with Mackes et al. (2018), a *more diverse* set of stimuli or longer video exposures might produce richer variance in observers’ annotations and potentially reveal additional empathic processes.
2. **Single-Cohort Observers:** Our sample primarily comprised university-aged participants. Broader demographic ranges (e.g., varying in age, cultural background, or clinical status) could yield different patterns of *annotation style* and empathic engagement, limiting the generalizability of our results.
3. **Annotation Constraints:** In *Experiment 2*, we restricted participants to discrete emotional categories, which might overshadow *fine-grained intensity* judgments. Future studies could deploy *hybrid rating scales* (e.g., time-continuous plus discrete labeling) to disentangle the interplay between classification and dynamic tracking.
4. **Physiological Focus:** While we examined target physiology (e.g., heart rate) to assess emotional intensity, we did not explore parallel psychophysiological measures from the observers. Investigating observer-specific responses (e.g., observer heart rate or skin conductance) could shed light on the *real-time* interactive dance of affective empathy—particularly whether certain participants exhibit stronger experience sharing under different annotation regimes.
5. **Potential for Social Desirability Bias:** As in many empathy paradigms, observers might have over-reported *perspective-taking* or certain empathic states. Incorporating *behavioral or neural* indices (e.g., fMRI or reaction time tasks) could reduce reliance on self-report measures and mitigate social desirability confounds.
6. **Limited Emotional Categories:** We primarily focused on certain discrete emotions (e.g., sadness, fear, happiness). Further research should investigate whether more complex or mixed-affect states (e.g., bittersweet joy or embarrassment tinged with humor) respond differently to time-continuous vs. discrete-annotation methods.

Taken as a whole, these considerations advocate for a *cautious interpretation* of our results and highlight directions in which this line of research can expand.

### 6.5 Future Directions

Our findings suggest that *continuous* emotion-intensity judgments may harness mentalizing capacity more robustly than *explicit categorical* labeling, which can disrupt perspective-taking correlations. Follow-up work could incorporate *neural imaging*, more *diverse participant samples*, and combined continuous–categorical scales to clarify the synergy between cognitive empathy (mentalizing) and affective empathy (resonance). Additionally, splitting broader emotional experiences into finer subcomponents—and measuring observer physiology—may help uncover whether discrete labeling tasks hamper or enhance certain empathic pathways. Ultimately, these refinements would inform not just empathy theory but also practical tools for *clinical assessment* and *intervention*, where harnessing or guiding different forms of empathic response can have substantial therapeutic implications.

In conclusion, *Experiment 1* successfully replicated and supported Mackes et al. (2018), illustrating how *time-continuous, open-ended* emotion ratings align with perspective-taking. *Experiment 2* revealed that *category-specific* annotation alters these correlates, paralleling arguments by Jospe et al. (2020) that methodology can recalibrate how observers engage cognitive vs. affective empathy. These dual experiments underscore empathy’s remarkable *flexibility*: individuals can swiftly shift between mentalizing and sharing, contingent on the instructions and rating schemes presented to them.

## AUTHOR CONTRIBUTIONS

AO conceived and designed the study. AO and SS designed the experimental stimuli. AO and SK developed the experimental procedure. AO, SK, AN, and SS collected the data. AO and SK performed the data analysis. AO created the visualizations and drafted the manuscript. SK and AM revised the manuscript critically for important intellectual content. All authors reviewed and approved the final version of the manuscript.

## ACKNOWLEDGMENTS

The authors would like to thank NTT ExCPartner Corporation for managing the overall production of the video materials used in this dataset. We also acknowledge the support of Intercross Corporation for acquiring physiological data during the video production and conducting Experiment 1, and NTT TechnoCross Corporation for implementing Experiment 2. Finally, we extend our sincere appreciation to all participants who took part in the study.

## SUPPLEMENTAL DATA

No supplemental material is available for this article.

## DECLARATION OF GENERATIVE AI TECHNOLOGIES IN THE WRITING PROCESS

During the preparation of this work, we used OpenAI’s ChatGPT (GPT-4 and GPT-5) to improve the clarity and readability of the manuscript’s language. After using these tools, we reviewed and edited the text as needed and take full responsibility for the content of this publication.

## DATA AVAILABILITY STATEMENT

The EMPAC dataset includes time-continuous emotion ratings, ECG and EDA signals collected from both professional actors and observers. Due to institutional and legal requirements, the dataset is currently available only to researchers affiliated with institutions in Japan. Data access is granted for academic purposes upon reasonable request and subject to review by the data access committee at NTT. Interested researchers may contact the corresponding author or for further details. We are continuing to assess possibilities for broader data sharing in compliance with ethical and legal standards. Information about the dataset is available in both English and Japanese on the official EMPAC project website: https://www.rd.ntt/e/dtc/gc1/research/empathy-dataset.html (English) / https://www.rd.ntt/dtc/gc1/research/empathy-dataset.html (Japanese).

